# Biomod2EZ - An R script suite for visualizing projected niche model ensembles and reporting statistical results

**DOI:** 10.1101/140855

**Authors:** Paul R. Sesink Clee, Stephen Woloszynek, Mary Katherine Gonder

**Affiliations:** Drexel University, Department of Biology – Philadelphia, PA; Drexel University, Department of Electrical and Computer Engineering – Philadelphia, PA

## Abstract

Biomod2EZ is a suite of R scripts to use in conjunction with the Biomod2 R package by Thuiller et al. (2016) that is used to create ensemble ecological niche models using up to 11 different modeling techniques. Biomod2EZ adds to the functionality of Biomod2 (Thuiller et al. 2016) by incorporating a report generation feature, detailed script annotation, and sample dataset/tutorial to ease the transition from ecological niche modeling using a Graphical User Interface to the coding environment of the R framework (R Development Core Team, 2016).

## Introduction

Ecological niche modeling involves predicting the distribution of a species as a function of underlying ecological patterns across a study region and has become in-creaseingly useful in studies involving biodiversity, conservation, and disease occurrence. Use of these models has increased dramatically in recent years with the growing availability of environmental map layers created from remotely sensed data. Still, the choice of which particular niche model to utilize remains a daunting task for most researchers. In addition to the variety of models available, one model may perform well for a specific dataset, but perform poorly with other models.

Biomod2 version 3.3-7 (Thuiller et al. 2016) is an R package that can currently be used to run up to 11 different niche model techniques on a single input dataset, calculates each model’s individual performance, and weights them accordingly to build an ensemble that combines that strength of each individual model. The contribution of models that perform poorly (determined by a user-defined threshold) can also be excluded from the ensemble. This ensemble technique has revolutionized the way that users can create ecological niche model projections without the need for in-depth knowledge of which single model is best suited for a given dataset. Although Biomod2 (Thuiller et al. 2016) is useful in creating ensemble models, it lacks a simple visualization of results that would be beneficial to users that are unfamiliar with R (R Development Core Team, 2016) and for model validation and interpretation.

## Functional Description and Advantages

Biomod2EZ is a suite of R scripts that aims to simplify the use of Biomod2 (Thuiller et al. 2016) for ecological niche modeling and automatically generate reports of model results. Many ecologists that are familiar with using Graphical User Interfaces (such as Maxent (Phillips et al. 2017)) are reluctant to create ecological niche models using the more complicated coding environment of the R framework. This script suite includes fully annotated scripts for users to input their data (i.e., species presence/absence, environmental layers, etc.) and create ensemble niche model projections that are exported directly into the working directory as an ASCII rasters for use in external mapping programs.

Biomod2EZ includes four scripts, an rmarkdown file for report generation, and a .jar file for Maxent (Phillips et al. 2017). The first script assists users in importing their presence/absence data, environmental predicting layers, and optional layers for projection (e.g., for projecting models to different geographic locations or under different climate scenarios). The second script is used to prepare the modeling environment and ensure that all associated packages are properly installed. The third script can be used to optionally remove duplicate presence and absence points that share the same coordinate. A final script contains all necessary steps to create niche models and a weighted ensemble along with modeling parameters that should be modified by users to best suit their datasets.

Most significantly, this R script suite improves the functionality of the Biomod2 package (Thuiller et al. 2016) by automatically generating reports that include output maps, performance scores for individual models, ensemble projections, decision trees for species presence/absence based on the provided environmental layers, environmental variable contribution to each individual model, indication of any failed models, and descriptions of individual models and test statistics used to determine model performance.

## Software Availability

This R script suite and included sample dataset/tutorial is available for download at: https://github.com/psesinkclee/biomod2ez

## Acknowledgements

Biomod2 is a product of Thuiller et al. (2016) and is implemented unaltered in this script suite to simplify use and generate a visual report of results.

